# Co-administration of midazolam and psilocybin: Differential effects on subjective quality versus memory of the psychedelic experience

**DOI:** 10.1101/2024.06.13.598878

**Authors:** Christopher R. Nicholas, Matthew I. Banks, Richard L. Lennertz, Cody J. Wenthur, Bryan M. Krause, Brady A. Riedner, Richard F. Smith, Paul R. Hutson, Christina J. Sauder, John D. Dunne, Leor Roseman, Charles L. Raison

**Affiliations:** Department of Family Medicine and Community Health, School of Medicine and Public Health, University of Wisconsin, Madison, WI 53706; Transdisciplinary Center for Research in Psychoactive Substances, University of Wisconsin, Madison, WI 53706; Department of Anesthesiology, School of Medicine and Public Health, University of Wisconsin, Madison, WI 53706; Department of Neuroscience, School of Medicine and Public Health, University of Wisconsin, Madison, WI 53706; School of Pharmacy, University of Wisconsin, Madison, WI 53706; Department of Psychiatry, School of Medicine and Public Health, University of Wisconsin, Madison, WI 53706; Wisconsin Institute for Sleep and Consciousness, University of Wisconsin, Madison, WI 53706; Center for Healthy Minds, University of Wisconsin, Madison, WI 53706; Department of Psychology, University of Exeter, Exeter, UK; Centre for Psychedelic Research, Imperial College London, UK

**Author notes:** <u>Corresponding authors:</u> Charles L. Raison, MD Department of Psychiatry, School of Medicine and Public Health University of Wisconsin-Madison 6001 Research Park Blvd Madison, WI 53719. <u>Corresponding authors:</u> Mathew I. Banks, PhD Department of Anesthesiology, School of Medicine and Public Health University of Wisconsin-Madison600 Highland Ave B6/319 Madison, WI 53792. Authors contributed equally to this paper (Christopher R. Nicholas, Matthew I. Banks).

## Abstract

Aspects of the acute experience induced by the serotonergic psychedelic psilocybin predict symptomatic relief in multiple psychiatric disorders and improved well-being in healthy participants, but whether these therapeutic effects are immediate or are based on memories of the experience is unclear. To examine this, we co-administered psilocybin (25 mg) with the amnestic benzodiazepine midazolam in 8 healthy participants and assayed the subjective quality of, and memory for, the dosing-day experience. We identified a midazolam dose that allowed a conscious psychedelic experience to occur while partially impairing memory for the experience. Furthermore, midazolam dose and memory impairment tended to associate inversely with salience, insight, and well-being induced by psilocybin. These data suggest a role for memory in therapeutically relevant behavioral effects occasioned by psilocybin. Because midazolam blocks memory by blocking cortical neural plasticity, it may also be useful for evaluating the contribution of the pro-neuroplastic properties of psychedelics to their therapeutic activity.

## Introduction

Serotonergic psychedelics such as psilocybin are associated with positive therapeutic outcomes in patients^1–11^ and enhanced well-being in healthy participants^12–14^, inducing durable benefit that far outlasts their acute pharmacological action. These benefits are assayed by validated scales of well-being^15–19^ and by narrative reports that describe the psychedelic experience as transformative even years later^12, 20, 21^. The efficacy of psilocybin administered with psychological support for the treatment of depression, anxiety, and substance use has driven a surge in interest in underlying mechanisms, especially the role of the acute experience versus memory for the experience.

Aspects of the experience induced by psilocybin, particularly those categorized as ‘mystical’, breakthrough, or insight-type, are predictive of long-term effects on well-being^18^ and of therapeutic benefit^3, 18, 22–28^. Inflexibility may be a common psychological trait underlying psychopathology^29, 30^; these acute experiences may promote psychological *flexibility* by promoting experiential engagement and openness to alternative beliefs about the self, others, and the world^28, 31^ and supporting change in the context of psychological and relational support integral to psychedelic-assisted therapy (PAT) protocols^32^. Thus, the transformative experience may facilitate immediate changes in perspective that correlate with positive outcomes.

Alternatively, or in addition, recapitulation of the experience after the dosing session (i.e., memory) may contribute causally to long-term behavioral change. The post-dose integration phase in which insights are translated into meaningful shifts in emotional awareness, narrative of self and world, and perspectives on behaviors may require long-term memory of the psychedelic experience to support therapeutic processing and long-lasting change^12, 33–35^. However, it is possible that to make sense of singular transformative experiences, study participants create *post-hoc* narratives that, while contributing to sense-making, do not carry any causal therapeutic power.

Testing these possibilities requires separating the transformative experience from its memory. We seek to do so using the benzodiazepine midazolam, an amnestic agent widely used in clinical settings. At low doses, midazolam is an amnestic agent^36^ that acts at GABAA receptors to block cortical neural plasticity^37–39^, the molecular correlate of memory. Consequently, midazolam may impair memory of the psychedelic experience while allowing the experience to occur. We sought to test this idea by co-administering midazolam and psilocybin. Because this has not been reported previously, we first determined the appropriate dose of midazolam in a pilot study. Here, we report the results of this dose-finding study. Our data suggest that midazolam can be safely co-administered with psilocybin, and that the ensuing psychedelic experience is of comparable subjective quality to that of psilocybin monotherapy. Importantly, the data suggest that midazolam impairs memory for the experience, potentially providing mechanistic insight into psilocybin’s therapeutic efficacy.

## METHODS

### Ethics statement

Research protocols were approved by the University of Wisconsin Institutional Review Board (Protocol #2020-0085); written informed consent was obtained from all participants. This study is registered on https://clinicaltrials.gov/study/NCT04842045.

#### Participants

Eight medically and psychiatrically healthy participants (4 female) were recruited via word of mouth, local advertisement, and www.clinicaltrials.gov. (See **Supplementary Methods** for more details.) After undergoing phone screening, medical and psychiatric eligibility assessments were conducted by a physician and psychologist, respectively. See **Supplementary Table 1** for full inclusion/exclusion criteria. See **Supplementary Table 2** for demographic data.

#### Study Design Overview

Psilocybin and midazolam were co-administered to all participants within a set and setting protocol that included preparatory sessions prior to dosing and integration sessions post-dosing (**Figure 1A**). On dosing day, participants were instrumented for scalp EEG recordings, administered midazolam and psilocybin, and assessed physically and psychologically (**Figure 1B**). The primary objective was to identify an optimal dosing strategy of midazolam that, when co-administered with an oral dose psilocybin (25 mg; Usona Institute, Fitchburg, WI), allowed a psychedelic experience to occur while impairing memory for that experience. Assessments were repeatedly conducted during the dosing session to assess the subjective quality of the psychedelic experience and to enable assessment of its memory post-dosing. Scalp EEG data were collected as objective measures of the effects of psilocybin. Memory for the experience and its personal salience were assayed on Day 1 and 8 post-dosing and well-being was assayed on Day 8 post-dosing. Adverse events were collected at all assessments. See **Figure 1A** for overall study scheme, and **Figure 1B** for dosing day schedule. See **Supplementary Methods** for further details, especially regarding preparatory and integration sessions.

**Figure 1.**
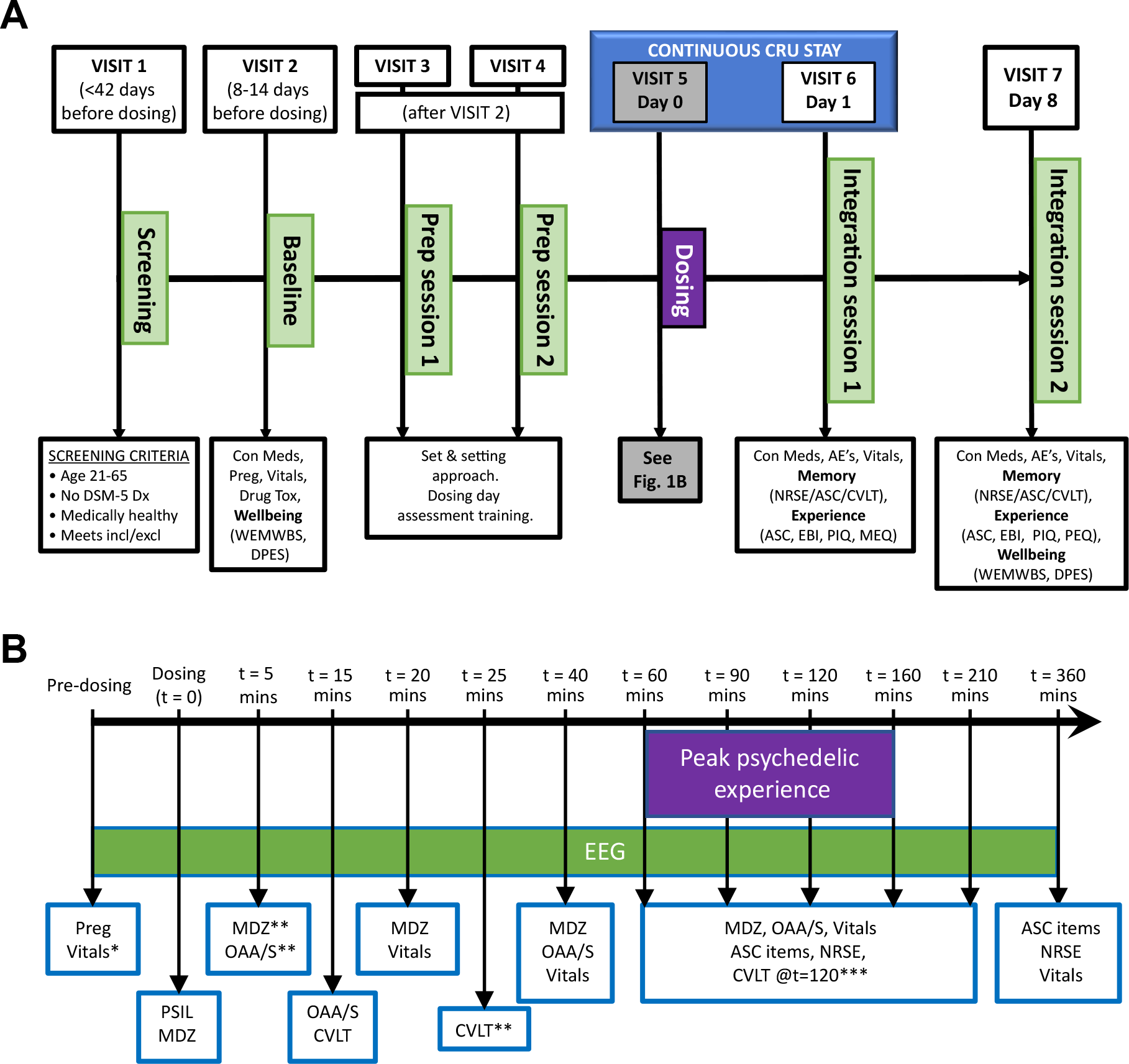
Study schema. A. Experimental schedule. AE = adverse event; ASC = Altered States of Consciousness questionnaire; CEQ = Challenging Experiences Questionnaire; Con Med = concomitant meds; CVLT = California Verbal Learning Task; DPES = Dispositional Positive Emotion Scales; Drug tox = urine drug screen; EBI = Emotional Breakthrough Inventory; MEQ = Mystical Experiences Questionnaire; NRSE = Narrative Report of Subjective Experience; PEQ = Persisting Effects Questionnaire; PIQ = Psychological Insight Questionnaire; Preg = pregnancy test; Vitals = blood pressure, heart rate, pO2; WEMWBS = Warwick Edinburgh Mental Well-being Scale. **B**. Dosing Day schedule. MDZ = midazolam; OAA/S = Observer’s Assessment of Arousal and Sedation; Preg = pregnancy test; PSIL = psilocybin. *Vitals = blood pressure, heart rate, pO2. **Participants 3-8 only. ***Participants 5-8 only

### Data collection and Analysis

#### Dosing Session

Dosing sessions occurred in the University of Wisconsin Hospital & Clinics Clinical Research Unit (CRU). Participants were supervised throughout by the facilitator(s), who also conducted preparatory and integration sessions with the participant. In addition to the facilitator(s), a board-certified anesthesiologist and a research coordinator accompanied the participant throughout the dosing session. After dosing, participants stayed overnight in the CRU, with discharge approximately 24 hours post-psilocybin dosing.

##### Midazolam administration

Midazolam is widely used for clinical sedation and has an excellent safety profile when administered according to established guidelines^40^. As an additional precaution, continuous pulse oximetry and supplemental oxygen were available at the direction of the anesthesiologist at all times. At appropriate doses, midazolam produces a state known as conscious amnesia^36^, in which participants are able to engage in conversation and cognitive tasks, but have little or no memory of the experience afterward^41^. This study targeted plasma concentrations (25 – 95 ng/ml) that produce conscious amnesia and incorporated graded dose adjustments based on functional assessment during the dosing session (**Supplementary Table 3 & 4**). The initial dose of midazolam was based on weight and age, and then adjusted by functional assessment during the dosing session (**Supplementary Table 3 & 4**). Following ingestion of psilocybin, an initial IV bolus of midazolam was administered followed by additional boluses over the next 3.5 hours, with the doses and timing based on a previously published pharmacokinetic model (**Figure 1B**)^42^.

Arousal level during the dosing session was assayed by the Observer’s Assessment of Arousal and Sedation (OAA/S)^43^. Scores on the OAA/S vary from 1 to 5, with 1 being completely unresponsive and 5 wide awake. An OAA/S score of 4 or 3 corresponds to mild or moderate sedation, respectively. Acute effects of midazolam on short term memory (unrelated to psychedelic experience content) were assessed via the California Verbal Learning Test (CVLT^44^), shown in previous reports^45, 46^ to be responsive to amnestic doses of midazolam. At t = 5 minutes after administration of psilocybin, the dose of midazolam was adjusted to target an OAA/S = 4. At t = 15 minutes, the CVLT was assessed, and the dose of midazolam was adjusted to target a CVLT score <25% that of baseline while still maintaining OAA/S ≧ 3. At t = 15 mins, 6/8 participants had OAA/S ≧ 4. At 25 and 120 minutes, the CVLT was administered to reassess the level of amnesia, but the dose of midazolam was not adjusted further based on the CVLT. Of the five OAA/S assessments between t = 40 and 210 mins, four participants always had OAA/S ≧ 4, three participants had one instance of OAA/S = 3, and one participant had two instances of OAA/S = 3. Although none of the participants had OAA/S scores < 3 at the designated assessment times, one participant became overly sedated between OAA/S assessments and their midazolam dose was reduced. To better impair memory, the midazolam dosing range was expanded considerably for participants 5-8 compared to the first four. This, combined with the dosing-day memory assessments, suggested that the cohort could reasonably be divided into two dosing groups (“high-dose” and “low-dose”) for descriptive purposes.

##### Dosing-day assessments

The subjective quality of the psychedelic experience was assessed in real time during the dosing period using selected questions from the Altered States of Consciousness (ASC) questionnaire^47^ (**Supplementary Table 5**), brief narrative reports, and, indirectly, using high-density EEG (hd-EEG; **Figure 1B**). These timing of these assessments was designed to capture the peak of the subjective experience, corresponding to the peak plasma concentrations of psilocin in our previous study^48^. The extent to which participants had a psychedelic experience was determined by comparing dosing-day ASC scores with normative data^26^. Because of our interest in the potential role of memory in mediating the persistent behavioral effects of psilocybin, we chose questions from the ASC previously observed to predict long-term antidepressant responses to psilocybin^26^. Specifically, ASC items #50, 77, 86, and 34 were the four most positively associated with therapeutic activity in that study, and item #85 was the most negatively associated. An additional question, “I saw colors in complete darkness or with closed eyes” was selected as a general check that a psychedelic-like experience was occurring. The questions are listed in **Supplementary Table 5**. Prior to answering these ASC items, participants provided 60-second free-form Narrative Reports of Subjective Experience (NRSE) that were audio-recorded. Five- to seven-word phrases were extracted from these reports to use for the post-dosing memory assessment. See below for procedure for obtaining these phrases.

#### Primary Objective and Endpoints

The primary endpoints were (1) the number of participants who had a psychedelic experience, and (2) the number of participants exhibiting memory impairment for that experience, when co-administered psilocybin and midazolam. These endpoints were assessed as follows:

1. To determine the occurrence of a psychedelic experience, the number of participants who, during the dosing session on Day 0, scored >50% of normative scores on selected questions from the ASC (**Supplementary Table 5**) was assessed. For each participant, ASC scores were calculated as the average of the maximum values on each question during the dosing session.
2. To determine the occurrence of memory impairment, the number of participants who, on post-dosing Day 1, scored <50% of the mean normative ASC scores was assessed.

Normative ASC scores are from a prior study involving psilocybin monotherapy for depression in which scores were obtained 24 hours after dosing^26^.

#### Secondary Objective and Endpoints

##### Post-dosing memory assessment

A secondary objective was to evaluate memory for the psychedelic experience following co-administration of midazolam and psilocybin using signal detection methods applied to ASC data. (Exploratory objectives involved applying the same methods to CVLT and NRSE data.) When assessed on Days 1 and 8 for memory of selected ASC items administered on dosing day, participants were directed to “rate to what extent the statements apply to the most intense portion of your particular experience during the dosing session – compared to normal waking consciousness” using the visual analogue scale on both Day1 and Day 8. Regarding yes-no recognition memory for previously encountering items and or foils, on Day 1 participants were instructed to answer, “specifically about your memory of previously seeing this statement and scale during your dosing session.” On day 8, participants were instructed to answer, “specifically about your memory of seeing each statement and scale at your previous follow-up visit.”

Data from this yes-no recognition protocol was used to calculate d’ (the difference between normalized hit rates and false alarm rates) as follows. (Exploratory objectives involved applying the same methods to CVLT and NRSE data.) On Day 1, participants were presented with items from the ASC and asked whether each item was old (i.e., from the dosing session) or new (i.e., foils). The questions were presented as Yes-No Recognition tasks with N=12 items, 6 old and 6 new. Hit-rates (H) and false alarm rates (F) were calculated as the proportion of hits and false alarms generated for old and new items respectively, using standard definitions^49^. To avoid generating infinite values for memory accuracy, H and F values of 0 and 1 were converted to 1 /(2N) and 1 – [1/(2N)], respectively, where N is the number of trials presented in the task (e.g. 0.042 and 0.958 for N=12). Memory accuracy was calculated as d’ = z(H) – z(F), where z(H) and z(F) are the z-values describing number of standard deviations from the mean that each hit and false alarm proportion represents on a normal distribution, respectively (e.g. z-value for 0.042 = -1.73; z-value for 0.5 = 0; z-value for 0.958 = 1.73). Overall, this approach generated a possible d’ range from 0 (chance levels – no recall) to 3.46 (no errors - perfect recall). Measurements were repeated on Day 8 to assess recovery of baseline memory function following elimination of psilocybin +/- midazolam. The Day 8 tasks included both items from Day 1 along with new distractors drawn from thematically similar scales not used on dosing day^50^.

##### Safety Assessment

An additional secondary outcome was to evaluate the safety of coadministration of psilocybin and midazolam. Adverse events were graded based on Common Terminology Criteria for Adverse Event Version 4.0. Adverse events were graded 1 - 5 depending on severity: grade 1 - mild, grade 2 - moderate, grade 3 - severe, grade 4 - life threatening, grade 5 - fatal.

#### Exploratory Objectives and Endpoints

Memory accuracy (d’) for CVLT and NRSE data was calculated as for the ASC. (See **Supplementary Methods** for more details.) Additional exploratory objectives were focused on examining associations between midazolam dosing, memory for the psychedelic experience, and psychological and behavioral assessments of salience, insight, and well-being. These included (at baseline and Day 8) the Warwick-Edinburgh Mental Well-Being Scale (WEMWBS) and the Dispositional Positive Emotion Scales (DPES), and (on Day 1) instruments previously associated with psilocybin effects on well-being^26, 51, 52^: (full) ASC, Mystical Experiences Questionnaire (MEQ), Emotional Breakthrough Inventory (EBI), and Psychological Insight Questionnaire (PIQ). We note that although the DPES is dispositional and therefore less likely to change, prior studies have demonstrated changes in dispositional traits after psilocybin dosing^24^. Long-term reflections on the psychedelic experience relevant to well-being were assessed on Day 8 using the final three questions of the Persisting Effects Questionnaire (PEQ)^53^; referred to as PEQ1 (“How personally meaningful was the experience?”), PEQ2 (“Indicate the degree to which the experience was spiritually significant to you.”), and PEQ3 (“Do you believe the experience led to change in well-being and life-satisfaction?”).

##### Analysis of resting state hd-EEG data

High-Density EEG data were recorded at a sampling frequency of 500 Hz with vertex-referencing, using a NetAmps 300 amplifier and NetStation software (Electrical Geodesics Inc., Eugene, OR). Standard preprocessing and spectral analysis were applied, as described^54^. Baseline recordings were obtained from participants (sitting quietly) prior to drug administration alternating between eyes closed and eyes open (4 minutes each condition). Power spectral densities (PSDs) from these recordings were compared to those computed from 10 minutes of data recorded ∼120 minutes after psilocybin administration, with analysis confined to data segments in which the participants had their eyes closed, and were not listening to music, being interviewed, or being presented with memory probe stimuli. See **Supplementary Methods** for further details.

#### Statistical Analysis

Because this was a dose-finding study, it was not powered to detect effects of midazolam or memory impairment on any of the outcome measures. Thus, the analyses presented here are solely for the purposes of generating hypotheses and should be interpreted with caution in the context of generalizing from these results. Linear regression analyses were applied to outcome measures as a function of midazolam dose and measures of memory, with the emphasis on the confidence intervals of regression slopes and adjusted coefficients of determination (*r*^2^).

## Data Availability

All data presented in figures in this paper will be made freely available at www.zenodo.com subject to constraints imposed by the University of Wisconsin IRB.

## Results

### Primary Endpoints

#### Midazolam dose-finding

A detailed description of the midazolam dosing strategy is presented in **Supplementary Tables 3** and **4**. The initial dose and number of dose adjustments during the dosing session increased during the study. For participants 1 & 2, the initial targeted plasma concentration of 25 ng/ml and a single increase to 35 ng/ml at 15 minutes did not produce adequate amnesia. For participants 3 & 4, the addition of a dose adjustment at 5 minutes and a final targeted plasma concentration of 45 ng/ml still did not produce adequate amnesia, as the participants still recalled >25% of words from the CVLT (**Figure 2A**). The initial targeted plasma concentration was then increased to 70 ng/ml with dose adjustments up to 125 ng/ml. Using this protocol, participants 5-8 all achieved criterion amnesia at targeted plasma concentrations between 65-95 ng/ml (**Figure 2A**) while exhibiting mild or moderate sedation (OAA/S scores 4 or 3, respectively; **Figure 2B**). Of note, some participants appeared to fall asleep for short periods of time in between assessments but awoke spontaneously or maintained arousal for 2-5 minutes after waking.

**Figure 2.**
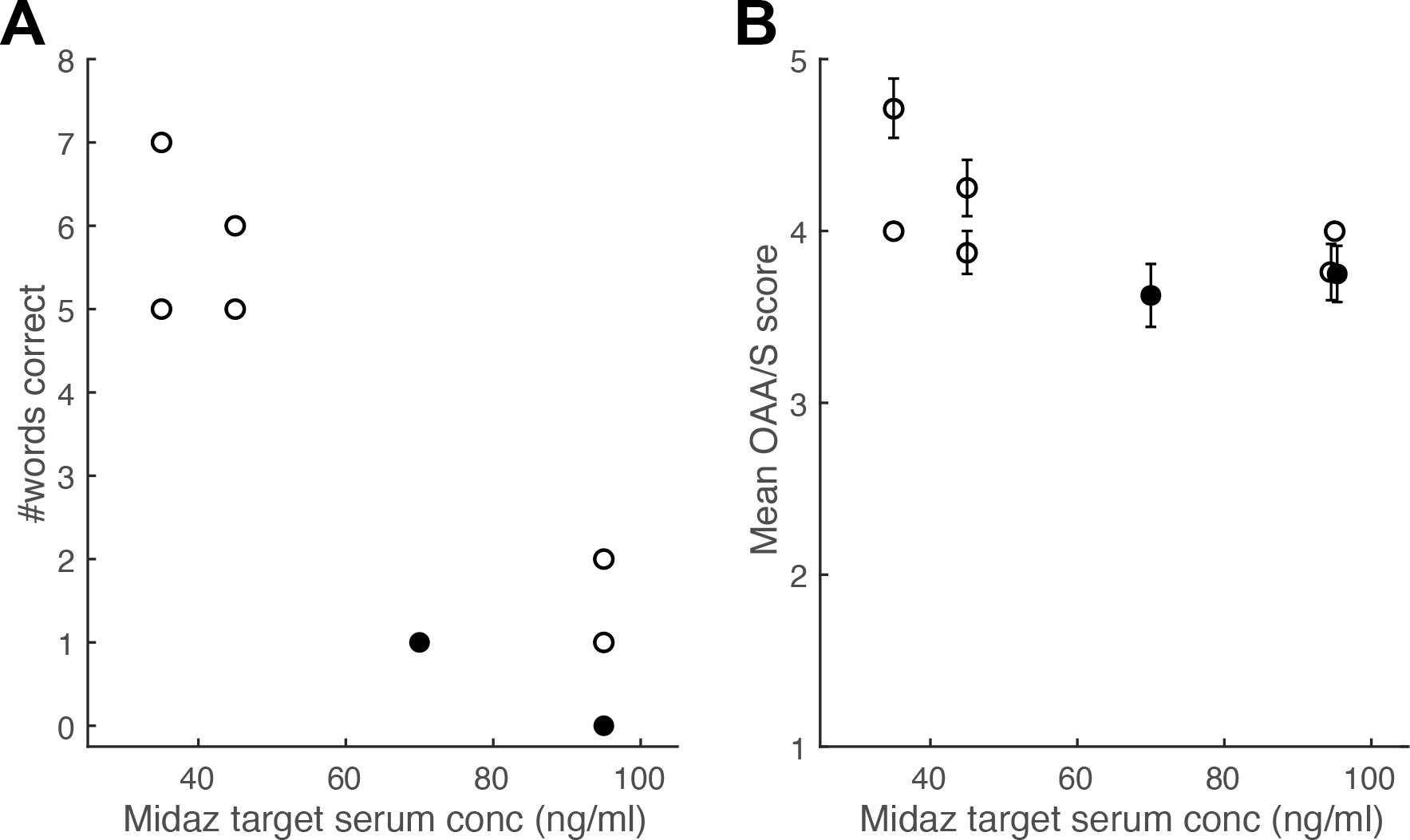
Dosing day assessments for the effects of midazolam. **A.** Performance on the California Verbal Learning Test (CVLT) administered 15 minutes after the initial midazolam bolus, as a function of midazolam (Midaz) target serum concentration. Each symbol represents one participant. Score range for the CVLT is 0 to 16. **B.** Observer’s Assessment of Arousal and Sedation (OAA/S) scores averaged over the dosing session as a function of midazolam target serum concentration. Each symbol is the OAA/S score for one participant averaged across the assessment times during the dosing session. Error bars represent standard deviation. The two participants who exhibited memory impairment in Figure 4 are indicated by *filled symbols*.

One participant briefly scored <3 on the OAA/S after increasing the targeted dose of midazolam. The dose was reduced according to **Supplementary Table 3**, after which the participant maintained an OAAS >3. Below, we refer to participants 1-4 as the “low-dose midazolam” group and participants 5-8 as the “high-dose” midazolam group. Overall, there was a strong association between dosing-day CVLT scores and midazolam dose, but not between OAA/S and midazolam dose (**Figure 2**; **Supplementary Table 6**).

#### Effect of midazolam on the acute psychedelic experience during its occurrence

To assess the subjective quality of the psychedelic experience, participants were repeatedly asked selected ASC questions during the dosing session. The time course of the subjective quality of the psychedelic experience was assessed by tracking mean scores on ASC questions during the dosing session (**Figure 3**). This time course was comparable to previous studies with psilocybin^55^, peaking about 2 hours after psilocybin administration (**Figure 3A**). Participants administered higher doses of midazolam did not experience a reduction in the magnitude or the duration of elevated ASC scores. Furthermore, most aspects of the experience exhibited similar trajectories on dosing day, though two, “I had particularly inventive ideas” and “time passed slowly in a painful way” were relatively less prominent in the experience compared to other ASC items (**Figure 3B**). Of note, the trajectories were as sustained—in fact were more sustained on average—for the high-dose midazolam group than for the low-dose group.

**Figure 3.**
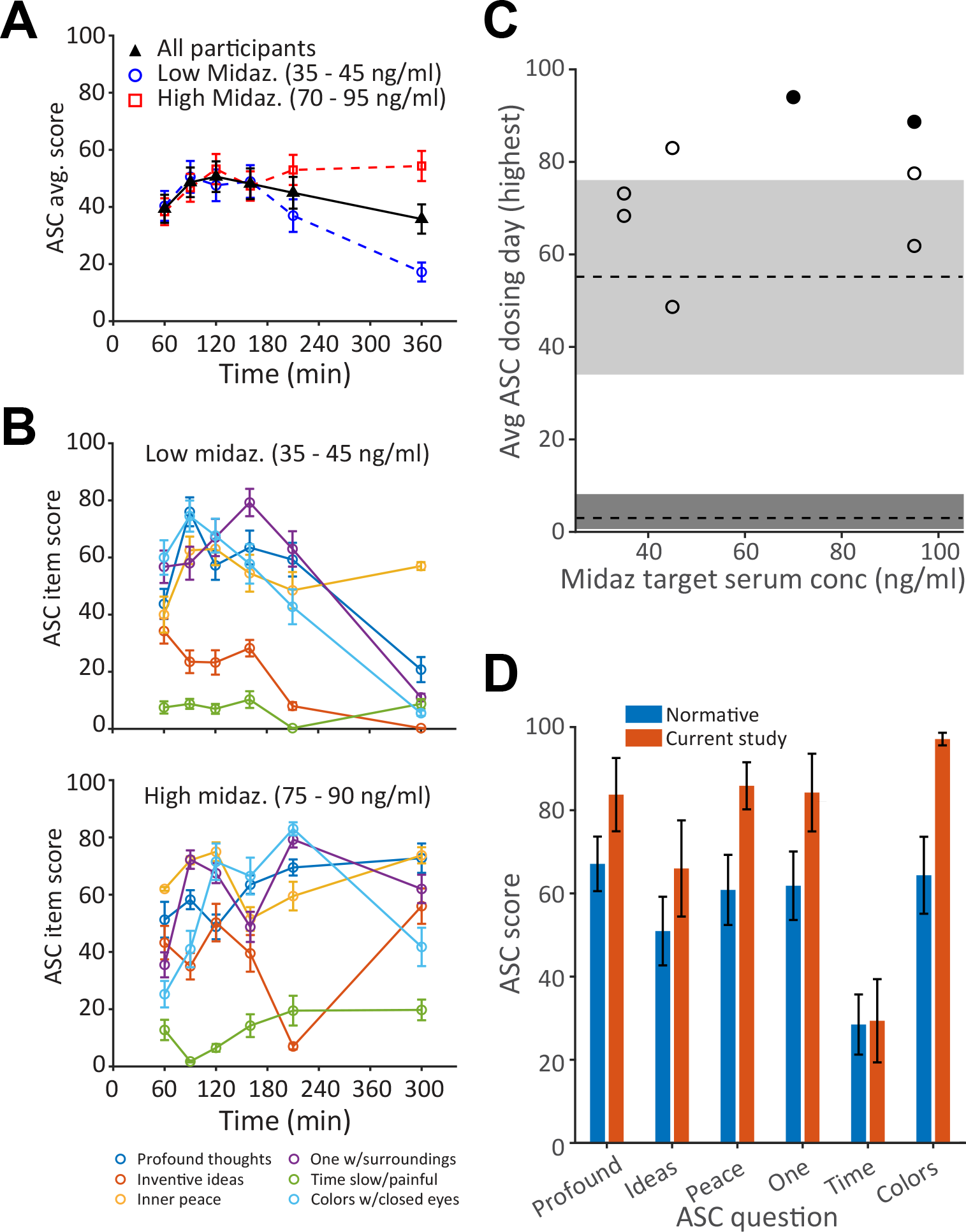
Evidence for maintained subjective quality of the psychedelic experience when psilocybin is administered with midazolam. A. Time course of subjective quality of the psychedelic experience, plotted as Altered States of Consciousness (ASC) questionnaire scores averaged across questions and participants. *Black symbols:* all participants. *Blue* and *red symbols:* low and high midazolam (Midaz) dose, respectively (n = 4 for each dose range). Score range for the ASC is 0 to 100. **B.** Scores on each question averaged across participants receiving low (*top*) and high (*bottom*) midazolam doses. **C**. ASC scores on dosing day for each participant as a function of midazolam dose. Symbols show average of the maximum score on each question during the dosing session for each participant. Grey rectangles represent median and interquartile range for normative data for psilocybin alone (median = 55)^26^ and for placebo (median = 3)^56^. **D.** Comparison of dosing-day responses on the ASC to normative data set^26^. Error bars display mean +/- SEM. See **Supplementary Table 5** for full text of ASC questions. The two participants who exhibited memory impairment in Figure 4 are indicated by *filled symbols*.

Consistent with the goal of maintaining the subjective quality of the psychedelic experience in the presence of midazolam, all participants scored >50% of the normative psilocybin ASC data during the dosing session, regardless of midazolam dose [**Figure 3C**; *grey rectangles* indicate normative ASC scores for psilocybin monotherapy (*upper rectangle*^26^) or placebo (*lower rectangle*^56^)]. In fact, participants tended to have higher scores than the normative data. Results were consistent across questions reflecting different domains of the psychedelic experience (**Figure 3D**).

#### Effect of midazolam on memory for the acute psychedelic experience

In contrast to the preserved psychedelic experience in the presence of midazolam, memory for the experience appeared to be impaired in some participants. None of the participants in the low-dose midazolam group met criteria for memory impairment. However, two of four participants in the high-dose midazolam group scored <50% of the mean normative ASC score on Day 1, one at a target midazolam serum concentration of 70 ng/ml, and one at 95 ng/ml (**Figure 4A)**.

**Figure 4.**
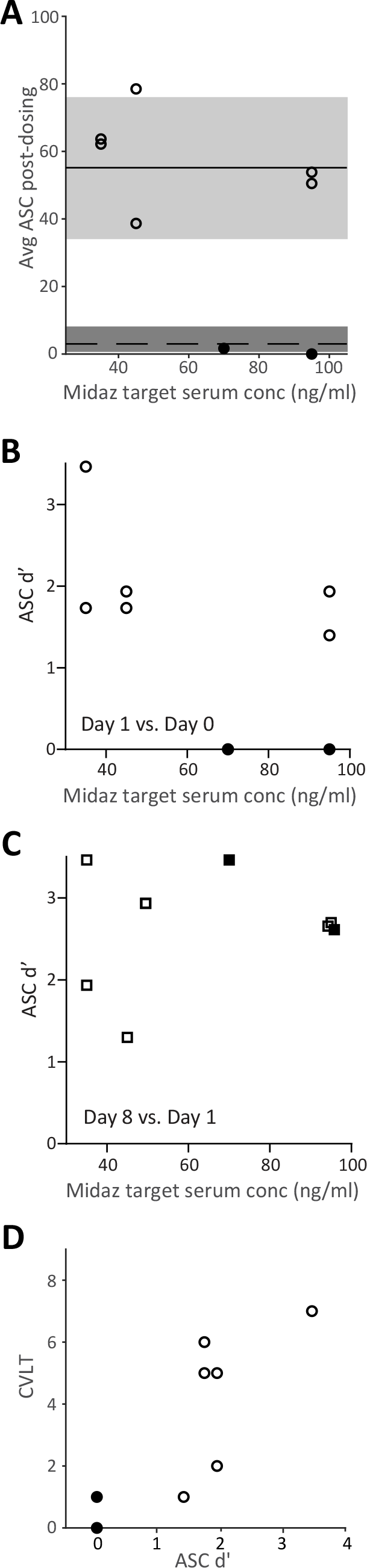
Evidence for impaired memory of the psychedelic experience when psilocybin is administered with midazolam. A. Altered States of Consciousness (ASC) questionnaire scores on Day 1 for each participant as a function of midazolam (Midaz) dose. Symbols show average of the scores on the 6 asked during the dosing session for each participant. (See **Supplementary Table 1**.) Grey rectangles represent median and interquartile range for normative data for psilocybin alone (median = 55)^26^ and for placebo (median = 3)^56^. **B.** Memory accuracy (d’) on day 1 for ASC items presented on dosing day. **C.** Memory accuracy (d’) on day 8 for ASC items presented on day 1 (*right*). **D.** CVLT performance as a function of dosing day memory accuracy (d’; Day 1 versus dosing day). *Filled symbols* indicate the two participants who exhibited memory impairment. Score range for d’ is 0 to 3.48.

### Secondary Endpoints

#### ASC item recognition accuracy

Recognition memory of the dosing day ASC items was used as a secondary assessment of memory of the psychedelic experience. Analysis of memory accuracy (d’; see Methods) for the dosing day yielded results consistent with the comparison of ASC scores to normative data presented in **Figure 4A**. Specifically, while none of the participants in the low-dose group exhibited evidence of impaired memory accuracy, the same two of four participants in the high-dose group who met the primary outcome criterion for ASC scores (**Figure 4B**) exhibited reduced d’ compared to the remaining participants. Furthermore, ASC d’ exhibited a modest tendency to decrease with increasing midazolam dose (**Supplementary Table 6**). Memory assessed on Day 8 for items presented on Day 1 did not demonstrate a trend with dose, indicating that memory impairment was specific for the dosing period (**Figure 4C**). Notably, there was a strong relationship between dosing-day memory assessments using the CVLT and measures of memory on Day 1 (**Figure 4D**, **Supplementary Table 6**), suggesting that dosing day amnesia is predictive of impaired memory post-dosing.

#### Safety of midazolam and psilocybin co-administration

The safety of psilocybin-midazolam co-administration was assessed by the incidence, type, and severity of adverse during and post-dosing. Six of eight participants experienced adverse events, including headache (n=4), migraine (n=1), jaw tightness (n=1), nausea (n=1), heavy limbs (n=1), back pain (n=1), urinary incontinence (n=1) emotional distress (n=1), and acid reflux (n=1). All events were mild to moderate in severity, and the majority resolved within 24 hours. Changes in heart rate and blood pressure during the dosing session were consistent with normative samples^57^ (**Supplementary** Figure 1).

### Exploratory Endpoints

#### CVLT and NRSE recognition memory

Recognition memory for the dosing day CVLT items and elements of the NRSE was analyzed on an exploratory basis. Memory accuracy (d’) for CVLT and NRSE items exhibited weak to modest trends toward decreasing values with increasing midazolam dose (**Supplementary** Figure 2AB; **Supplementary Table 6**).

#### Effects on insight and emotional salience

Effects of psilocybin on measures of emotional salience and personal insight, including the Emotional Breakthrough Inventory (EBI), Psychological Insight Questionnaire (PIQ), and summary questions from the Persisting Effects Questionnaire (PEQ), associate with reduction in depression symptoms^51, 52^. Here, we investigated whether midazolam tended to reduce psilocybin-induced long-term behavioral changes by investigating scores on these measures assessed on Day 1 (EBI, PIQ) or Day 8 (PEQ). We observed a trend for modest negative associations in the relationship between these measures and midazolam dose (**Supplementary** Figure 3A; **Supplementary Table 6**) and a trend for modest positive associations between all measures and memory of the psychedelic experience, as assayed by ASC d’ on Day 1 (**Supplementary** Figure 3B; **Supplementary Table 6**).

#### Effects on well-being

Similar to measures of salience and insight, measures of well-being associate with therapeutic activity in clinical trials with psychedelics^58^. Here, effects on well-being were assessed using the change in Warwick-Edinburgh Mental Well-Being Scale (WEMWBS^15^) and the Dispositional Positive Emotion Scales (DPES^59^) at baseline compared to Day 8. Across all participants, treatment effects on both measures were modest, with WEMWBS increasing by 0.75 ± 8.7 (mean ± SD; median = 1, range = - 15 to 17) and DPES increasing by 0.33 ± 0.28 (mean ± SD; median = 0.32, range = -0.05 to 0.71). These modest changes may have been due to a ceiling effect, as participant scores tended to be high even before psilocybin administration (e.g., for WEMWBS: median 56.5, range 40 – 69: all but one participant scored higher than the normative median = 51^15^). Nevertheless, change in well-being tended to decrease with increasing midazolam dose, with a stronger trend for DPES (**Supplementary Table 6**). In addition, when we compared change in WEMWBS to memory for the psychedelic experience, expressed as ASC d’ on Day 1 (cf. **Figure 4B**), we observed a trend for an association **Figure 5A**; **Supplementary Table 6**). However, no such trends were detected for change in DPES (**Figure 5B**; **Supplementary Table 6**). These data offer some suggestion that memory impairment may predict poorer long-term behavioral change following psilocybin administration.

**Figure 5.**
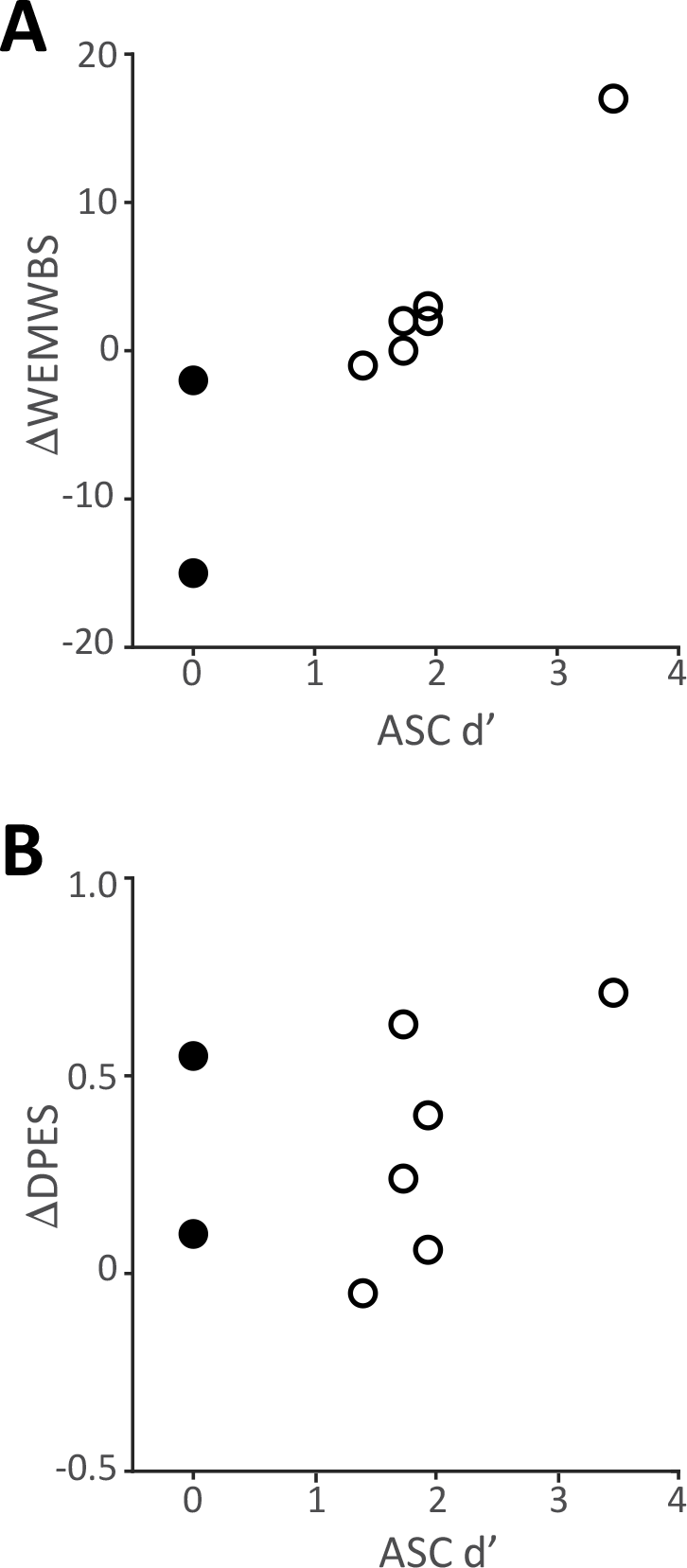
**Relationship between change in well-being and memory of the psychedelic experience**. Change in Warwick Edinburgh Mental Well-being Scale (**A**, ≧ WEMSBS) and change in Dispositional Positive Emotion Scales (**B**, ≧ DPES), measured as value on Day 8 minus baseline, is plotted as a function of ASC memory accuracy (d’; *right*). Each symbol represents one participant. Note that *smaller* d’ indicates *greater* memory impairment. The two participants who exhibited memory impairment in Figure 4 are indicated by *filled symbols*.

#### Effects on brain activity

Resting state scalp EEG data were obtained from participants during the dosing session. In control conditions, alpha band (8-12 Hz) power increases when participants close their eyes, an effect linked to visual cortex entering an ‘idling mode’ during periods of decreased visual information processing^60^. Psilocybin and other serotonergic psychedelics attenuate this increase in eyes-closed EEG alpha power^61–64^, likely because vivid visual hallucinations activate visual cortical areas^65^. We observed similar suppressive effects of psilocybin on eyes-closed alpha power on average (**Figure 6A**) and in 7 of 8 individual participants (**Figure 6B**), and this effect appeared to be independent of midazolam dose (**Figure 6B**, **Supplementary Table 6**). Topographic analysis of the distribution of alpha power across the electrode array confirmed that pre-treatment eyes-closed alpha power was focused in the back of the brain over visual cortex and was profoundly suppressed during psilocybin treatment (**Figure 6C**).

**Figure 6.**
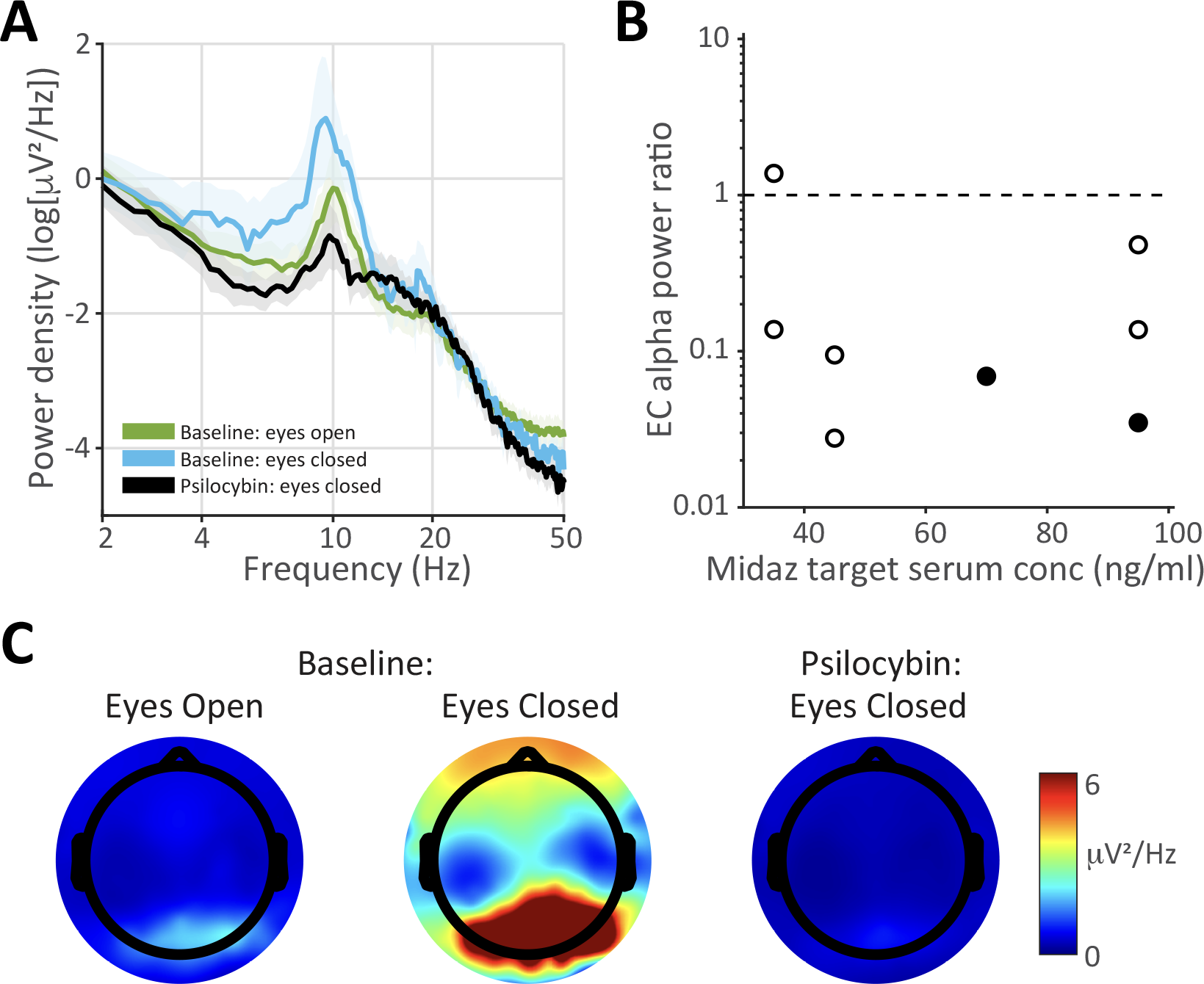
Eyes-closed (EC) electroencephalography (EEG) alpha power is suppressed by psilocybin in the presence of midazolam. A. Power spectrum at electrode Pz averaged across all subjects during baseline (pre-drug) eyes-open, baseline eyes-closed, and eyes-closed at t = 2 hrs. after psilocybin administration. Note the prominent alpha peak (8-12 Hz) during pre-drug eyes-closed that is suppressed during pre-drug eyes-open and during eyes-closed with psilocybin. Data clouds represent standard errors of the mean. **B.** Eyes-closed alpha power ratio (psilocybin/baseline) for each participant as a function of target midazolam (Midaz) dose. **C.** Distribution of alpha power across the EEG electrode array showing that eyes-closed alpha during baseline (*center*) is focused in the back of the brain and is profoundly suppressed at t = 2 hrs. after psilocybin administration. The two participants who exhibited memory impairment in Figure 4 are indicated by *filled symbols*.

## Discussion

There is considerable interest in identifying the mechanisms underlying the therapeutic activity of serotonergic psychedelics, given accumulating evidence of positive outcomes in both clinical and healthy samples.^11, 13, 22, 33, 66^. The relationship of the subjective quality of the psychedelic experience with clinical outcomes^23, 35, 67^ and evidence of pro-neuroplastic effects in rodent models^68, 69^ suggest that the therapeutically-relevant effects of psychedelics reflect a complex interplay between subjective effects, especially salience and insight, and cellular and molecular effects. Within a standard PAT model, the post-dose integration phase requires that memory for the psychedelic experience be accessible and intact to harness the drug’s therapeutic benefit^34, 70, 71^. In this pilot study, we used the co-administration of midazolam with psilocybin to conduct an initial exploration of the mechanistic role of acute experiential and pharmacological effects versus memory for the psychedelic experience. Midazolam is a benzodiazepine that induces “conscious sedation” and amnesia, permitting acute conscious experiences while suppressing the memory for these experiences. Here, we identified a dosing strategy for midazolam that when co-administered with psilocybin produced minimal sedation and allowed the occurrence of an acute psychedelic experience comparable to normative data but that appeared sufficient to attenuate memory of the psychedelic experience. Identifying this dosing strategy is a prerequisite for a subsequent placebo-controlled trial examining the role of memory and conscious experience in therapeutic outcomes in patient populations treated with psilocybin.

The dosing day evaluation of the acute psychedelic experience using selected questions from the ASC showed that all participants exhibited a psychedelic experience in the presence of midazolam comparable to normative ASC psilocybin data without midazolam. Five of 6 ASC items presented during the dosing session were selected based on strong associations with therapeutic benefit in a previous study involving psilocybin monotherapy^26^. All participants had average scores on these ASC items that were >50% of the normative data comparison^26^, as specified in the study design, an effect that appeared to be independent of midazolam dose. Indeed, 7 of 8 participants rated the subjective quality of the experience higher than the normative median, suggesting that midazolam might enhance the perceived quality of the psychedelic experience during its occurrence at the doses administered here.

Maintenance of the subjective quality of the acute psychedelic experience during midazolam is also supported by exploratory analysis of eyes-closed EEG alpha power, which was reduced in 7 of 8 participants independent of midazolam dose. Suppressed alpha power has previously been reported for serotonergic psychedelics^62–64^, likely due to the prevalence of visual hallucinations^65^. These data indicate that key features of the psychedelic experience were preserved in the presence of midazolam.

The effects on memory were more subtle. Although dosing day CVLT scores indicated profound amnestic effects of midazolam in the high dose group, only 2 participants had post-dosing ASC scores <50% of the normative data, the specified study endpoint for memory. However, secondary analyses of dosing-day ASC items suggest an inverse relationship between midazolam dose and memory accuracy, consistent with impaired memory. Complementary exploratory analyses of dosing-day CVLT items and narrative elements yielded weaker effects but were also consistent with memory impairment.

Safety data indicate that the co-administration of midazolam and psilocybin was safe and well-tolerated. Modest elevations in blood pressure were observed, presumably in response to psilocybin administration, as reported previously^53, 72^.

Post-dosing memory assessments indicated partial memory impairment but not complete amnesia. This contrasts with the pronounced amnestic effects of these midazolam doses in previous studies^36^. It is possible that the experiential activation, i.e., the profound alterations in sensory and cognitive processing, induced by psilocybin in the current study, competed against the amnestic properties of midazolam observed in previous studies.

Memory of the psychedelic experience relies on (likely cortical) neural plasticity induced during the acute experience. Based on its amnestic effects and its ability to block cortical neural plasticity^37–39^, midazolam may be a pharmacological tool to disambiguate the mechanistic contribution of neural plasticity compared to the psychedelic experience in the therapeutic effects of psychedelics. This is relevant given the central role proposed for neural plasticity in the mechanisms of psychiatric disorders, especially in key brain regions including prefrontal cortex^73–76^. Blunted neural plasticity is a presumed biological substrate of being “stuck” in a constricted worldview with “rigid priors,” i.e., psychological inflexibility^77^, and has been described as a transdiagnostic factor^30, 78^ with regards to symptom severity and expression in psychiatric disorders, and more broadly to diminished flourishing and negativity bias^73, 79–81^. Although purely speculative, midazolam may also block cortical neural plasticity in areas such as the prefrontal cortex that are postulated to contribute to the therapeutic effects of psilocybin. There are several important caveats to this idea. First, the plasticity underlying memory and the plasticity underlying persistent behavioral effects of psilocybin are likely distinct, though possibly overlapping.

Second, although midazolam blocks plasticity outside of hippocampus^39^, its effects in prefrontal cortex are untested. Third, benzodiazepines not only *block* cortical neural plasticity, but they can also *induce* subcortical neural plasticity, e.g., in dopaminergic circuits^82, 83^, which may form the basis for their addictive potential. Testing this idea would require a follow-up study in which neural plasticity is measured directly during administration of psilocybin +/- midazolam.

Psilocybin also affects memory^84^, likely via its actions on serotonin 5HT2A receptors in areas of the brain responsible for memory encoding and retrieval. This may be the mechanism whereby these drugs promote post-dosing psychological flexibility^85^. In preclinical models, serotonergic psychedelics modulate behavioral phenotypes and induce rapid (within hours) and long-lasting (weeks) neural plasticity^68, 69, 86–88^, paralleling the time course of rapid symptomatic relief in patients^11, 23, 58, 66^. Acutely induced changes in gene expression may alter circuits in prefrontal cortex and connected regions controlling emotional processing^89, 90^. These changes may facilitate subsequent behavioral changes during the post-dosing psychotherapeutic “integration” sessions that are standard practice in clinical trials. A challenge in determining the relative contribution of experience versus plasticity to psilocybin’s long-term behavioral effects is that their dose ranges entirely overlap, and thus it is difficult to separate them experimentally. Indeed, the interaction between the two may be important, and thus both may contribute^91, 92^. Midazolam may help disentangle these different mechanisms.

Serotonergic psychedelics induce a biochemical stress response in preclinical models^93^ and human participants^57^, as well as fear- and anxiety-producing experiences^26^. There is uncertainty about the potential contribution of these stressful components of the experience to its therapeutic benefit^26, 94, 95^. Acute stress also modulates memory formation, adaptively promoting memory of the stressor and suppressing memory of peripheral elements of the experience^96^. Benzodiazepines are anxiolytic at doses even lower than those causing amnesia (e.g., <40 ng/ml serum conc.)^46, 97^, and thus are expected to suppress the psilocybin-induced biochemical stress response and any stressful components of the psychedelic experience. In the current study, decreased stress may have contributed to the observation that ASC scores on dosing day were high relative to normative data and remained elevated for longer during the dosing session for high-dose compared to low-dose midazolam. It is also possible that midazolam’s effects on the stress response contributed indirectly to impaired memory for the experience on Day 1 and Day 8.

The observed effects of psilocybin administered with midazolam on well-being were more modest than those reported in response to psilocybin alone^98, 99^. It is likely that this was due at least in part to a ceiling effect, as baseline scores on the WEMWBS in the current study were already elevated compared to normative data for these measures^15^. However, we did observe a trend for WEMWBS to decrease with increasing memory impairment. The absence of comparable effects on DPES may be due to the latter measure capturing only an aspect of well-being, i.e., positive emotions related to self and others, although an inverse association was observed between midazolam dose and DPES score. In addition, in contrast to the partial effect of midazolam on memory of the psychedelic experience, the effects on measures of insight and emotional salience (EBI, PIQ, PEQ) were more dramatic. These data suggest that even partial blockade of memory blunts the long-term behavioral effects of psilocybin. Alternatively, it is also possible that midazolam acutely suppressed insight and emotional salience during the dosing session, and that these aspects of the psychedelic experience were not captured by the selected ASC items administered during dosing. That is, the effect of midazolam on post-dose measures of psychological insight, emotional breakthrough, and meaningfulness may have been secondary to an acute effect on these features during the psychedelic experience. If so, this acute effect may also contribute to the amnestic effects of midazolam, given the dependence of memory on emotion^100^.

Indeed, both the acute stress response induced by serotonergic psychedelics^57, 72^ and the emotional and personal salience of the experience may modulate memory formation^100, 101^, likely through effects on attention^102^. Midazolam would be predicted to suppress these memory-supporting mechanisms, raising the possibility that it could be used experimentally to distinguish between specific subjective features of the psychedelic experience (e.g. higher-order personal salience vs hallucinogenic effects).

### Limitations and future directions

The small number of participants limits formal significance testing of the relationships between memory, EEG, and behavioral assays and midazolam dose. Because it was a dose-finding study, there was also no placebo control or assessment of variables such as expectation^103^, suggestibility^104^, therapeutic alliance^51^, and blinding effectiveness^105^. The normative ASC data used for comparison were obtained from a patient population 24 hours after psilocybin administration, rather than from healthy volunteers during the dosing session. A planned follow-up study with placebo controls will enable more rigorous quantitative analysis, formal hypothesis testing, and assessment of these variables. Further, there was no evaluation of insight and emotional salience of the psychedelic experience during the dosing session; inclusion of assays in these domains in future studies will elucidate the contribution of insight and salience to memory formation and help distinguish direct from indirect effects of midazolam on these domains. Intermittent somnolence was also observed in 4 participants during the dosing session, even though participant reports supported the occurrence of a psychedelic experience comparable to normative data. This suggests that arousal level and/or connection to the environment may dissociate from the psychedelic experience in this context, but the current study included no formal assessment of these effects. The methodology of collecting and analyzing the NRSE in the current study is not yet fully developed and will require reliability testing in future studies. Regarding the memory assessments, there may have been a more pronounced practice effect with the ASC items than with the CVLT or NRSE items, due to the ASC items being presented multiple times throughout the dosing session at regular intervals. Reduced repetition or single use of ASC items unique to the memory battery could address this in future work, though the trade-off is reduced resolution of potentially relevant subjective effects across the dosing session. Additionally, recognition of the NRSE items may have been inflated due to recognition of non-episodic content cues, such as the presence in the foils of idiosyncratic semantic terms a participant would consider themselves unlikely to use^106^. Assessment for recall of experiential content prior to assessing recognition of NRSE items would provide an alternative approach to assess the effects of cue distinctiveness on memory facilitation. Probing recall and recognition of audio and visual stimuli would provide additional dimensions for assessing alterations in episodic memory. Use of metacognitive efficiency measures and^107^ independence remember/know models to assess changes to memory using dual process signal detection^108, 109^ are also planned for future work, once larger sample sizes are available to support these approaches.

## Conclusions

In this dose-finding study, we identified a dose of the amnestic agent midazolam that, when co-administered with psilocybin, allowed a psychedelic experience to occur of comparable intensity and subjective quality to psilocybin alone, but impaired memory for the experience. The dose of midazolam and extent of memory impairment tended to associate inversely with salience, insight, and changes in well-being. These data suggest the potential role of memory in the persistent behavioral effects induced by psilocybin. Furthermore, although midazolam did not appear to dampen the subjective psychedelic experience based on our current measures, previous reports suggest that it blocks memory by blocking cortical neural plasticity. The roles of cortical neural plasticity versus the subjective psychedelic experience are a matter of active debate^110, 111^. A fully powered placebo-controlled follow-up study will enable a more rigorous investigation of the role of experience, memory, and neural plasticity in the persistent behavioral effects of serotonergic psychedelics.

## Acknowledgements

Funding provided by Mary Sue and Mike Shannon and the UW-Madison Department of Anesthesiology. We are grateful to Kristin Van Hyfte and Lucy Ptak for serving as research coordinators, Katie Chilton, MD for acting as a facilitator, to Michael Sutherland for assistance with EEG data analysis, and for the time and effort of our participants.

