## Supplementary information for "Co-administration of midazolam and psilocybin: Differential effects on subjective quality versus memory of the psychedelic experience"

[*] Authors contributed equally to this paper

### SUPPLEMENTARY METHODS

#### Psilocybin source and safety considerations.

Psilocybin was obtained as 25 mg capsules from Usona Institute (Fitchburg, WI). Psilocybin can produce psychologically meaningful yet challenging experiences for healthy participants. However, numerous studies over the last 25 years have shown that psilocybin is safe when administered in a controlled, psychologically supportive setting[1]. In the current study, the research team had prior experience administering psychedelic compounds in the context of a clinical trial[2, 3].

#### Participants

Eligible participants were medically and psychiatrically healthy, non-smoking adults between the ages of 18 and 65 years-old who had not used a psychedelic substance in >3 months. Participants with a body mass index >35, a blood pressure >140/90, or significant abnormalities on physical exam or laboratory studies were excluded from dosing. Similarly, participants with a major *Diagnostic and Statistical Manual of Mental Disorders*, fifth edition (DSM-5) diagnosis based on the Mini International Neuropsychiatric Interview (MINI[4]) or a family history of psychosis or bipolar disorder based on the Family History Screen (FHS[5, 6]) were excluded from dosing. Eligible participants were also required to perform within age-based normative limits on the CVLT.

#### Preparatory Sessions.

Participants received up to four hours of pre-dose preparation with a clinical psychologist facilitator (and when available, a co-facilitator) over two visits. At least one visit occurred in a private room within the University of Wisconsin Hospital and Clinics Clinical Research Unit (UWHC-CRU) to familiarize the participant to the dosing session environment. If needed, one of the preparation sessions could occur virtually using a HIPAA compliant video platform. Preparation sessions adhered to a “set-and-setting” approach[7] focused on building rapport, reviewing important aspects of the participants life-history, setting intentions and expectations for their participation in the study, and education about study and safety procedures. To avoid contamination of the post-dosing memory assessments by the Day 1 and Day 8 integration sessions, participants were informed that the facilitator was not permitted to discuss the content of the experimental stimuli and assessment measures, observations, or verbal content shared by the participant about their dosing session experience until completion of the Day 8 memory assessment. During the preparation sessions, participants were provided training on the dosing session assessments, which included review of the narrative assessment procedure and practice responding to mock visual-analog stimuli unrelated to items on the ASC administered on the dosing day.

#### Integration sessions.

Day 1 and 8 integration sessions occurred after completion of all memory and self-report assessments. The purpose of the integration sessions was to provide an opportunity for the participant to discuss any remembered elements of their dosing session experience, including any subjective content from the session that may have affected their thoughts, emotions, and behaviors. Facilitator(s) took a supportive, non-directive approach and refrained from confirming or interpreting the participants’ experiences, rather only reflecting back any insights or observations by the participant that might assist with integrating their experience into their everyday life. Upon completion of the final Day 8 memory assessment and self-report measures, the facilitator was permitted to discuss any content or observations about which the participant was interested in learning and answer any questions about the dosing session experience.

**Analysis of resting state hd-EEG data**.

High-Density EEG (HdEEG) data were recorded at a sampling frequency of 500 Hz with vertex-referencing, using a NetAmps 300 amplifier and NetStation software (Electrical Geodesics Inc., Eugene, OR). After applying a first-order high-pass filter (single pole IIR, 0.1 Hz) to eliminate direct current shifts, data were band-pass filtered (2-way least squares FIR, 1–50 Hz) using Matlab software (The MathWorks Inc., Natick, MA). As with other studies using 256-channel sensor arrays, electrodes on the face and outer ring of the sensor net were removed leaving 185 channels[8]. Additional noisy channels that were consistently out of range of baseline based upon visual inspection were also removed. On average, 98% or more of the 185 channels were retained. Using EEGLAB[9] we then visually inspected and removed sections of noisy data samples that contained obvious movements or large areas of artifact spanning multiple channels before running the data through EEGLAB's Adaptive Mixture ICA (AMICA) to decompose the data into individual components. The data was then inspected again and time periods with poor decomposability, where noise still contaminated multiple components, were removed from the data and AMICA was repeated on the original data with these time periods removed. Note, multiple AMICAs were run only to identify bad data segments. For the final data, only one AMICA was computed on the clean data. Components containing eye, muscle, and channel artifact were then removed. Missing channels were then interpolated using EEGLAB's spline interpolation and average referenced. Spectral power density was computed using Welch's modified periodogram method (*pwelch* function in MATLAB) in 2-s Hamming windows (50% overlap) to decompose EEG time series signals into frequency bands of interest.

### Analysis of Narrative Report of Subjective Experience (NRSE)

All phrases were structured to express a single, complete idea. All constructed phrases began with a verb focused on the action or perception of the participant; verbs for constructed phrases were matched with verbs for distractor items presented in the yes-no recognition tasks administered post dosing (see below). The NRSE and questionnaire items were grouped together with assessments of vital signs to limit the number of interruptions to the psychedelic experience.

#### Phrase Generation Protocol:

*To select the phrases, first identify the time point that has the highest total score on the 11-item 5D-ASC measurement; _______ minutes. This is the 'peak' time point.*

*Nine phrases should be generated from the narrative across this 'peak' time point and the time points most closely preceding and following the 'peak' time point.*

*Wherever possible, three phrases should be generated from the peak time point, three should be generated from time points occurring before the peak, and three should be generated from time points occurring after the peak.*

*Phrases should be between five and seven words long. Phrases should express a single complete idea, and should be centered on action / perception of the subject. The implicit subject of each phrase should be "I". Wherever possible, the verbs 'Saw, Explored, Realized, Heard, Encountered, Met' should be used to start the phrase. (i.e. Met all of my past selves)*

*Alternative preferred verbs to select from would include 'Felt, Touched, Tasted, Smelled, Understood, Recognized, Became, Left, Transformed, Grew, Shrank, Gave, Received'.*

*Within the nine generated phrases, there should ideally be three paired phrases that share the same verb. (i.e. Realized the universe is made of love ; Realized my mother had forgiven me)*

*Six phrases (including three of the paired phrases) will then be presented to the subject alongside the distractor items on post-dosing day 1.*
*The three remaining paired phrases will be presented with the non-paired phrases on post-dosing day 8. (i.e. Phrases presented on post-dosing day 1 would include both 'met all of my past selves' and 'realized the universe is made of love', while post-dosing day 8 phrases would include both 'met all of my past selves' and 'realized my mother had forgiven me'.)*
*This will ensure that half of the phrases presented at memory battery on post-dosing day 8 are novel, while half of the phrases have been previously encountered.*

*Finally, the order of presentation for the generated phrases and distractors should be randomized during the memory battery (as with all memory battery items from the CVLT).*

*Phrase 1: ______________________________________________________ (Unique Verb – Present on post-Dosing Days 1 and 8)   Phrase taken from Time Point: ________ min*

*Phrase 2: ______________________________________________________ (Unique Verb – Present on post-Dosing Days 1 and 8)   Phrase taken from Time Point: ________ min*

*Phrase 3: ______________________________________________________ (Unique Verb – Present on post-Dosing Days 1 and 8)   Phrase taken from Time Point: ________ min*

*Phrase 4: ______________________________________________________ (Verb matched with 7 – Present on post-Dosing Day 1 )   Phrase taken from Time Point: ________ min*

*Phrase 5: ______________________________________________________ (Verb matched with 8 – Present on post-Dosing Day 1)   Phrase taken from Time Point: ________ min*

*Phrase 6: ______________________________________________________ (Verb matched with 9 – Present on post-Dosing Day 1)   Phrase taken from Time Point: ________ min*

*Phrase 7: ______________________________________________________ (Verb matched with 4 – Present on post-Dosing Day 8)   Phrase taken from Time Point: ________ min*

*Phrase 8: ______________________________________________________ (Verb matched with 5 – Present on post-Dosing Day 8)   Phrase taken from Time Point: ________ min*

*Phrase 9: ______________________________________________________ (Verb matched with 6 – Present on post-Dosing Day 8)   Phrase taken from Time Point: ________ min*

**Supplementary Table 1. Inclusion and exclusion criteria**

| |
|---|
| **Inclusion Criteria** |
| 1. Individuals aged 18 – 65 years. |
| 2. Willing and able to provide informed consent. |
| 3. Willing to comply with all study procedures and be available for the duration of the study. |
| 4. Able to read, speak and understand English. |
| 5. Ability to take oral medication. |
| 6. Warwick Edinburgh Mental Wellbeing Scale (WEMWBS) score less than or equal to 53. |
| *7.* Women of childbearing potential must use an acceptable method of contraception from 30 days prior to dosing and agree to use an acceptable method of contraception until study completion. Males must agree to avoid impregnation of women throughout study participation through use of an acceptable method of contraception. *Note: Includes, but is not limited to, barrier with additional spermicidal foam or jelly, intrauterine device, hormonal contraception (started at least 30 days prior to study enrollment), intercourse with men who underwent vasectomy.* |
| 8. Eastern Cooperative Oncology Group (ECOG) performance status of 0 based on physical exam, which indicates that the participant is fully functioning. |
| 9. Agree to refrain from using legal psychoactive substance for the following defined time periods (the exception is caffeine): |
| · Tobacco and Nicotine: from Screening until Study Termination |
| · Alcohol: 72 hour window prior to the Dosing Visit |
| 10. Non-smokers. |
| 11. No psychedelic substance use within 3 months prior to their investigational product dosing visit by self-report. |
| **Exclusion Criteria** |
| 1. Calculated Body Mass Index (BMI) >35 at Screening |
| 2. Clinically significant abnormal chemistry or hematologic laboratory results (from a screen of Complete Blood Count with Differential and Comprehensive Metabolic Panel and urinalysis), using the UWHC lab reference intervals. |
| 3. Urine drug test containing non-prescribed drugs of abuse (i.e., non-prescribed opioids, benzodiazepines, amphetamines, cocaine) at Baseline. |
| 4. Evidence of ischemic disease, ventricular arrhythmias, or cardiac conduction defects on electrocardiogram (ECG) at Screening. |

| |
|---|
| 5. History of Prolonged QT syndrome as determined by self-report or QTc interval > 450 milliseconds as calculated by the Screening ECG. |
| 6. Clinically significant abnormalities on physical examination. |
| 7. Women of childbearing potential with positive urine pregnancy test at Screening or Dosing or woman who are breastfeeding. |
| 8. At Screening and Baseline, pre-hypertension defined as systolic blood pressure ≥140/90 and tachycardia/bradycardia defined as heart rate ≥ 100 beats per minute or < 45 beats per minute. On dosing day, acute hypertension/tachycardia/bradycardia (systolic blood pressure ≥ 140 mmHg or diastolic blood pressure ≥ 90 mmHg, heart rate > 100 beats per minute or < 45 beats per minute). |
| 9. Currently meets diagnostic criteria for any DSM-5 psychiatric condition as assessed by the MINI International Neuropsychiatric Interview (MINI). Prior history of primary psychotic disorder (unless substance-induced or due to a medical condition), bipolar disorder Type I or Type II, or schizophrenia, as determined by the MINI will be exclusionary. |
| 10. Concurrent or recent (within 1 year) history of major depressive disorder, obsessive-compulsive disorder, generalized anxiety disorder, panic disorder, anorexia nervosa, or bulimia nervosa, posttraumatic stress disorder, as determined by the MINI and psychiatric history. |
| 11. Active suicidal ideation or attempt in prior 12 months as assessed by the MINI and/or Columbia Suicide Severity Rating Scale (C-SSRS) |
| 12. First degree family history of primary psychotic disorder, bipolar disorder Type I, bipolar disorder Type II, or schizophrenia, as determined via self-report. |
| 13. MRI-incompatible implants or devices such as certain cardiac pacemakers or defibrillators, insulin pumps, cochlear implants, metal in orbit, implanted neural stimulators, central nervous system (CNS) aneurysm clips and other medical implants that have not been certified for functional magnetic resonance imaging (MRI). |
| 14. Not able to fit in the scanner coil (e.g., weight greater than 300 pounds, very large shoulders, etc.) |
| 15. Use of psychotropic medications, including antidepressants, mood stabilizers (lithium, carbamazepine, valproic acid, lamotrigine), antipsychotics, benzodiazepines, psychostimulants, or supplement-type antidepressants/anxiolytics (St. John's Wort, SAMe, L-methylfolate, valerian) within 3 months of Screening. |
| 16. Any regular use of medication, with the exception of females taking birth control or individuals taking medications approved for use by the study PI (or designee). |
| 17. As determined by self-report, history of cardiovascular disease, stroke, seizure disorder, acute narrow angle glaucoma, clinically-significant autoimmune conditions, claustrophobia, neoplasm (other than resected basal or squamous cell skin cancer), Type I or insulin-dependent Type II diabetes. |
| 18. Chronic viral infection, including hepatitis C, hepatitis B and human immunodeficiency virus |

| |
|---|
| 19. Inability to perform California Verbal Learning Test (CVLT), assessed by screening CVLT score of less than or equal to 5th percentile. |
| 20. Lack of a local support person who is available during the participant's 24-hour treatment and observation period, as determined by self-report. |
| 21. No intravenous (IV) access by self-report based on prior attempts. |
| 22. Any physical or psychological symptom, based on the clinical judgment of the study physician and/or psychologist that would make a participant unsuitable for the study. |

**Supplementary Table 2. Demographic data for study participants.**

| Participant | Age | Sex | Ethnic identity | Racial identity |
|---|---|---|---|---|
| 1 | 30 | M | Non-Hispanic | White |
| 2 | 57 | F | Non-Hispanic | White |
| 3 | 28 | M | Non-Hispanic | White |
| 4 | 42 | F | Non-Hispanic | White |
| 5 | 26 | F | Non-Hispanic | White |
| 6 | 59 | M | Hispanic | Unknown |
| 7 | 31 | F | Non-Hispanic | White |
| 8 | 58 | M | Non-Hispanic | White |

**Supplementary Table 3: Midazolam dosing algorithm**

| Assessment Timepoint | Scale(s) | Metric | Midazolam Dosing |
|---|---|---|---|
| 0 minutes | N/A | N/A | 44 mcg/kg bolus (adjusted by age). Start at Level 2. |
| 5 (±3) minutes post psilocybin/initial midazolam dosing | OAA/S | OAA/S = 1-2 | See **Note** below for guidance. |
| | | OAA/S = 3 | Maintain midazolam dosing at Level 2. |
| | | OAA/S = 4-5 | Increase midazolam dosing by 1 level. |
| 15 (±5) minutes post psilocybin/initial midazolam dosing | OAA/S and CVLT | CVLT <25% and OAA/S = 1-2 | See **Note** below for guidance. |
| | | OAA/S = 3<br>or<br>CVLT <25% & OAA/S = 4-5 | Maintain midazolam dosing. |
| | | CVLT ≥25% & OAA/S = 4-5 | Increase midazolam dosing by 1 level. |
| 25 (±5) and 210 (±10) minutes post psilocybin/initial midazolam dosing | CVLT | N/A | The CVLT at 25 (±5) minutes will be used to verify level of memory impairment after dose adjustments and prior to the psychedelic experience. The CVLT at 120 (±10) minutes will be used to verify level of memory impairment during the peak psychedelic experience. No dose adjustment should be made based on the CVLT at these time points. |
| 40 (±10), 60 (±10), 90 (±10), 120 (±10), 160 (±10), 210 (±10) minutes post psilocybin/initial midazolam dosing | OAA/S | N/A | Dose level may be adjusted to prevent over-sedation per the study physician's discretion. See **Note** below for guidance if OAA/S score reaches 1 or 2. |
| **Note:** If a subject becomes sedated to an **OAAS of 2** at any time, their midazolam dosing tier will be decreased. This does not apply to a subject who has fallen asleep (see below). If a subject becomes over-sedated to an **OAA/S of 1**, no further midazolam will be administered. The subject will be placed on continuous pulse oximetry. Supplemental oxygen and flumazenil will be administered at the discretion of the study physician. This does not apply to a subject who has fallen asleep. If the study physician judges that a participant has fallen asleep, they will wake the subject and reassess sedation in 2-5 minutes. Continued sedation to an OAAS of 1 will be considered oversedation. | | | |

OAA/S = Observer's Assessment of Arousal and Sedation; mcg/kg = micrograms per kilogram; CVLT = California Verbal Learning Test

**Supplementary Table 4: Midazolam administration**

| | Midazolam dose (mcg/kg) | | | | | | | | |
|---|---|---|---|---|---|---|---|---|---|
| Time (minutes) | 0 | 5 | 20 | 40 | 60 | 90 | 120 | 160 | 210 |
| **Level 1** | 44 | 0 | 0 | 22 | 15 | 15 | 15 | 15 | 15 |
| ***Level 2** | 44 | 0% | 22 | 22 | 22 | 22 | 22 | 22 | 22 |
| **Level 3a** | 44 | 31 | 22 | 31 | 31 | 29 | 29 | 29 | 29 |
| **Level 3b** | 44 | 0 | 44 | 31 | 31 | 29 | 29 | 29 | 29 |
| **Level 4** | 44 | 70% | 31 | 40 | 40 | 40 | 40 | 40 | 40 |

Intermittent boluses of midazolam (mcg/kg) were administered to participants for 3.5 hours following psilocybin administration. The doses above are presumed to be given to a 40-year-old participant and was adjusted by 1% for every year older/younger than 40 years, such that older patients received less medication than younger patients. The initial bolus (at time 0) was based on total body weight. Subsequent doses were based on adjusted body weight (ideal body weight + [total body weight - ideal body weight]*0.4). Dosing started at Level 2 and was adjusted according to level of sedation (Observer's Assessment of Arousal and Sedation [OAA/S]) and recall (California Verbal Learning Test [CVLT]) as outlined in **Supplementary Table 3**. Cells with bold borders illustrate the doses received by an example study participant who moved from dosing level 2 to dosing level 3b at 20 minutes based on their CVLT assessment.

**Supplementary Table 5**. Dosing-day Altered State of Consciousness (ASC) questionnaire items.

| ASC item# | Prompt |
|---|---|
| 50 | I had particularly profound thoughts |
| 77 | I had particularly inventive ideas |
| 86 | I experienced a profound inner peace |
| 34 | I felt one with my surroundings |
| 85 | Time passed slowly in a painful way |
| 22 | I saw colors in complete darkness or with closed eyes |

**Supplementary Table 6. Linear regression fit parameters.**

| Figure | dependent var | independent var | slope | slope CI | Adj $r^2$ | $t$ | $p$ |
|---|---|---|---|---|---|---|---|
| Fig. 2A | $CVLT_{Dosing}$ | Mdz | -0.0880 | [-0.128, -0.0476] | 0.797 | -5.33 | 0.00178 |
| Fig. 2B | OAAS | Mdz | -0.0166 | [-0.0439, 0.0108] | 0.267 | -1.68 | 0.168 |
| Fig. 3C | $ASC_{Dosing}$ | Mdz | 0.151 | [-0.363, 0.665] | -0.074 | 0.719 | 0.499 |
| Fig. 4A | $ASC_{Day1}$ | Mdz | -0.519 | [-1.42, 0.386] | 0.1216 | -1.40 | 0.210 |
| Fig. 4B | ASC d' | Mdz | -0.0228 | [-0.0566, 0.0109] | 0.200 | -1.66 | 0.149 |
| Fig. 4D | $CVLT_{Dosing}$ | ASC d' | 1.93 | [0.550, 3.31] | 0.605 | 3.42 | 0.0141 |
| Supp. Fig. 2A | CVLT d' | Mdz | -0.0153 | [-0.0404, 0.00976] | 0.150 | -1.49 | 0.186 |
| Supp. Fig. 2B | NRSE d' | Mdz | -0.0177 | [-0.0381, 0.00278] | 0.331 | -2.11 | 0.0789 |
| | MEQ | Mdz | -0.0238 | [-0.0664, 0.0188] | 0.110 | -1.36 | 0.221 |
| Supp. Fig. 3A, top left | EBI | Mdz | -0.982 | [-1.67, -0.293] | 0.615 | -3.49 | 0.0130 |
| Supp. Fig. 3A, top right | PIQ | Mdz | -0.0474 | [-0.0819, -0.0129] | 0.596 | -3.36 | 0.0152 |
| Supp. Fig. 3A, bottom left | PEQ1 | Mdz | -0.0449 | [-0.104, 0.0145] | 0.257 | -1.85 | 0.114 |
| | PEQ2 | Mdz | -0.0391 | [-0.0772, -0.00102] | 0.431 | -2.51 | 0.0457 |
| Supp. Fig. 3A, bottom right | PEQ3 | Mdz | -0.0346 | [-0.0637, -0.00564] | 0.519 | -2.92 | 0.0265 |
| | ∆WEMWBS | Mdz | -0.137 | [-0.422,0.147] | 0.0539 | -1.18 | 0.282 |
| | ∆DPES | Mdz | -0.00851 | [-0.0143, -0.00270] | 0.628 | -3.58 | 0.0116 |
| | MEQ | ASC d' | 0.783 | [-0.122, 1.69] | 0.332 | 2.12 | 0.0785 |
| Supp. Fig. 4A, top left | EBI | ASC d' | 15.0 | [-10.2, 40.3] | 0.138 | 1.45 | 0.196 |
| Supp. Fig. 4A, top right | PIQ | ASC d' | 0.922 | [-0.182, 2.03] | 0.312 | 2.04 | 0.0870 |
| Supp. Fig. 4A, bottom left | PEQ1 | ASC d' | 0.746 | [-0.919, 2.41] | 0.0282 | 1.10 | 0.315 |
| | PEQ2 | ASC d' | 0.718 | [-0.411, 1.85] | 0.169 | 1.56 | 0.171 |
| Supp. Fig. 4A, bottom right | PEQ3 | ASC d' | 0.844 | [0.126, 1.56] | 0.510 | 2.88 | 0.0282 |
| Fig. 5A | ∆WEMWBS | ASC d' | 6.95 | [3.52, 10.4] | 0.771 | 4.96 | 0.00256 |
| Fig. 5B | DPES | ASC d' | 0.0861 | [-0.152,0.324] | -0.0316 | 0.886 | 0.409 |
| Fig. 6B | EC $\alpha$ ratio | Mdz | -0.005 | [-0.0209, 0.0110] | -0.0631 | -0.765 | 0.474 |

Slope CI = 90% confidence intervals; Adj $r^2$ = adjusted coefficient of determination; $t$ = $t$-statistic; $p$ = uncorrected $p$-values. CVLT = California Verbal Learning Test; midaz = midazolam; OAAS = Observer's Assessment of Arousal and Sedation; ASC = Altered States of Consciousness Questionnaire; NRSE = Narrative Report of Subjective Experience; MEQ = Mystical Experiences Questionnaire; EBI = Emotional Breakthrough Inventory; PIQ = Psychological Insight Questionnaire; PEQ = Persisting Effects Questionnaire; WEMWBS = Warwick Edinburgh Mental Wellbeing Scale; DPES = Dispositional Positive Emotion Scales; EC = eyes closed

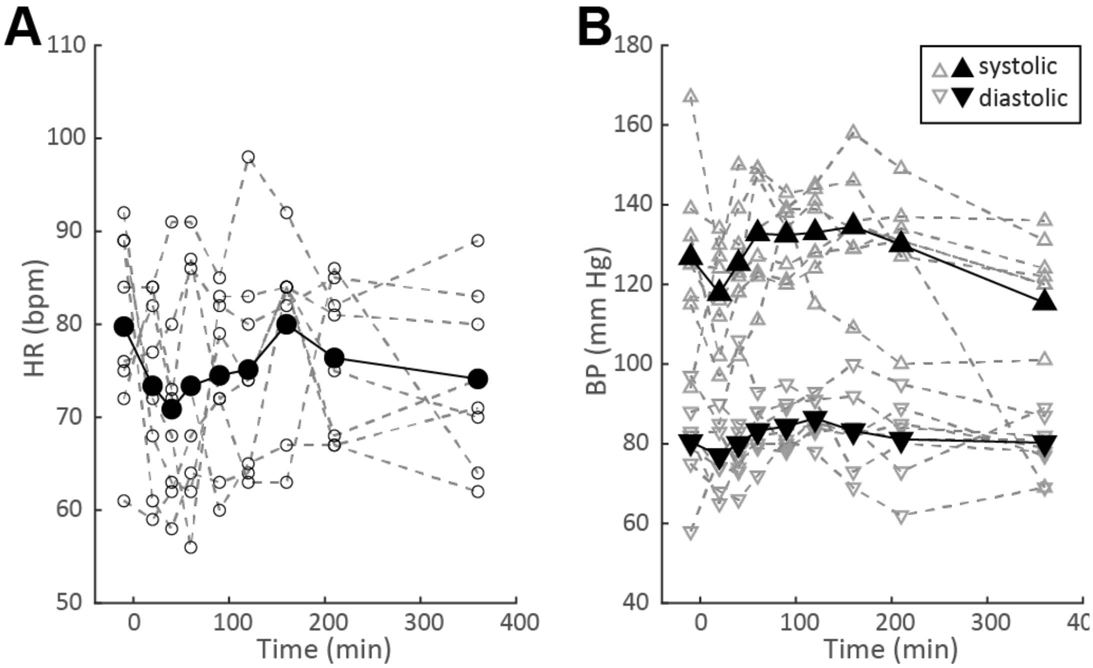


**Supplementary Figure 1. Dosing day physiological measurements.** Data are plotted as a function of time relative to psilocybin administration. *Small grey symbols: individual participants. Large black symbols: mean across participants.* **(A)** Heart rate (HR). **(B)** Blood pressure (BP).

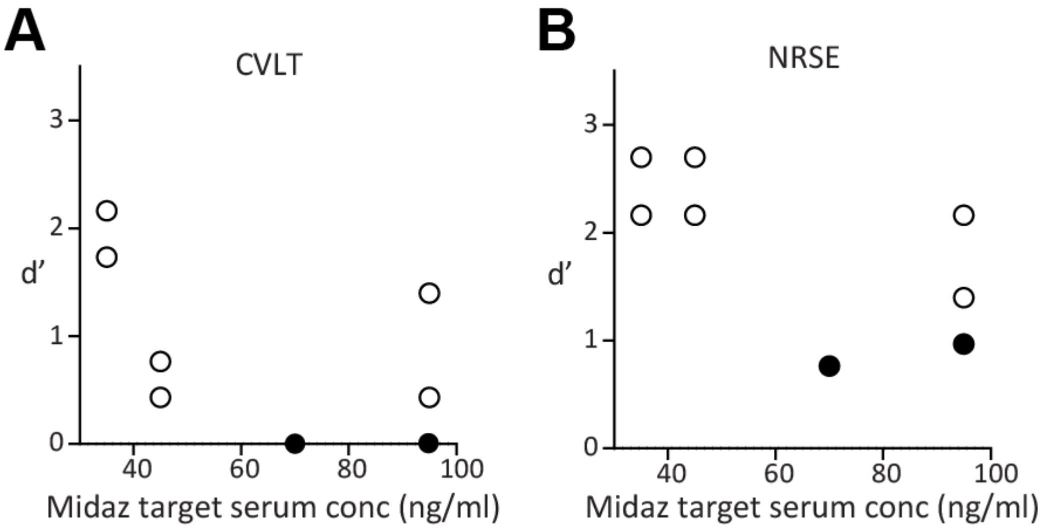


**Supplementary Figure 2. Memory accuracy (d') for California Verbal Learning Test (CVLT; A) and Narrative Report of Subjective Experience (NRSE; B) as a function of midazolam (Midaz) dose**. d' measured on Day 1 relative to dosing day responses. Each symbol is a participant. Score range for d' is 0 to 3.48.

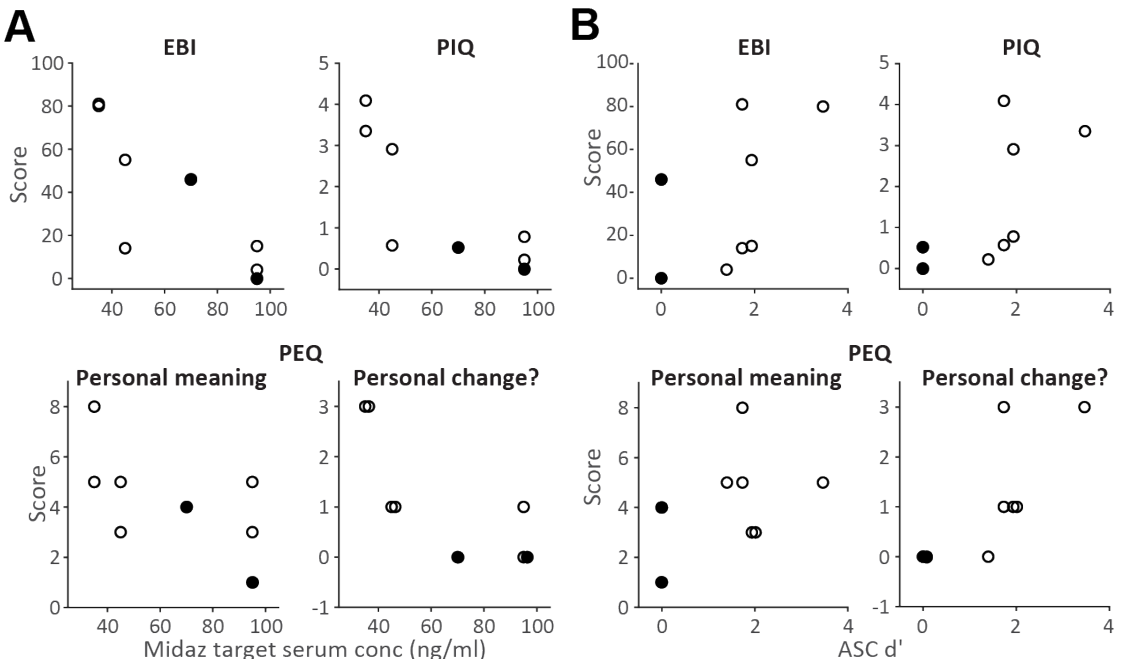


**Supplementary Figure 3. Effects on emotional salience.** Scores on questionnaires evaluated salience are plotted as a function of midazolam (Midaz) dose (**A**) and as a function of memory accuracy (d') for the Altered States of Consciousness (ASC) questionnaire items asked on dosing day (**B**). Each symbol represents one participant. EBI = Emotional Breakthrough Inventory (score range = 0 to 100); PIQ = Psychological Insight Questionnaire (score range = 0 to 5); PEQ = Persisting Effects Questionnaire summary questions ("How personally meaningful was the experience?", score range = 1 to 8; "Do you believe the experience led to change in well-being and life-satisfaction?", score range = -3 to 3).
